# TissUUmaps 3: Improvements in interactive visualization, exploration, and quality assessment of large-scale spatial omics data

**DOI:** 10.1101/2022.01.28.478131

**Authors:** Nicolas Pielawski, Axel Andersson, Christophe Avenel, Andrea Behanova, Eduard Chelebian, Anna Klemm, Fredrik Nysjö, Leslie Solorzano, Carolina Wählby

**Affiliations:** Department of Information Technology and SciLifeLab BioImage Informatics Facility, Uppsala University, Uppsala, 752 37 Sweden

**Keywords:** Interactive visualization, spatial omics, spatial transcriptomics

## Abstract

**Background and Objectives:** Spatially resolved techniques for exploring the molecular landscape of tissue samples, such as spatial transcriptomics, often result in millions of data points and images too large to view on a regular desktop computer, limiting the possibilities in visual interactive data exploration. TissUUmaps is a free, open-source browser-based tool for GPU-accelerated visualization and interactive exploration of 10^7+^ data points overlaying tissue samples.

**Methods:** Herein we describe how TissUUmaps 3 provides instant multiresolution image viewing and can be customized, shared, and also integrated into Jupyter Notebooks. We introduce new modules where users can visualize markers and regions, explore spatial statistics, perform quantitative analyses of tissue morphology, and assess the quality of decoding in situ transcriptomics data.

**Results:** We show that thanks to targeted optimizations the time and cost associated with interactive data exploration were reduced, enabling TissUUmaps 3 to handle the scale of today’s spatial transcriptomics methods.

**Conclusion:** TissUUmaps 3 provides significantly improved performance for large multiplex datasets as compared to previous versions. We envision TissUUmaps to contribute to broader dissemination and flexible sharing of large-scale spatial omics data.

## 1 Introduction

Over the past few years, a variety of spatial transcriptomic (ST) techniques has emerged [47], leading to scientific advances in many different areas[6]. These techniques can be classified into two groups: imaging-based ST (IST), which directly sequences individual mRNA molecules [26, 27, 31, 61, 62, 18, 22], and sequencing-based ST (SST), which uses micro-arrays or fluorescent beads to capture and sequence mRNA molecules with maintained information on spatial location [55, 50, 56]. The techniques have their respective advantages and disadvantages, but they share a common difficulty: the vast number of targeted molecules and the wide range of spatial scales makes visual exploration difficult. For example, a single IST experiment can produce millions of spatial markers for various mRNA molecules. Molecular patterns on a micrometer scale define cell types while the centimeter scale is needed for identifying tissue-level structures. However, patterns appearing at different scales are difficult to interpret using only one resolution and instead require interactive multi-resolution viewing. Moreover, interactive visualization of large quantities of markers is computationally challenging. Sub-sampling information, as is common in map-viewing functions such as Google Maps, could be an option, but may be risky: Sub-sampling sparse signals representing the presence of e.g. a virus, or a rare cell type, may lead to loss of very relevant information.

Additionally, the large number of different molecules targeted in parallel poses practical and computational challenges when it comes to interactive visualization. A single IST experiment can target up to hundreds of different mRNA species. Exploring patterns in such high-dimensional spaces is fundamentally difficult. A common approach to make the data more manageable is to compute the composition of molecules within small regional areas, such as segmented cells or spatial bins, to generate high-dimensional feature vectors [45, 48, 42, 41]. These features can then be projected into a lower-dimensional feature space using techniques such as UMAP [10] or t-SNE [59] to reveal different cell types or spatial domains. Unlike single-cell RNA sequencing data, ST data includes information about the location of RNAs within the imaged tissue. To fully make use of this information, visualization must simultaneously show the detected RNAs in their spatial context as well as their location in feature space. Here, region-based markers, such as polygons, can be used to visualize segmented cells. Regions may also be manually or automatically drawn to represent, e.g., tumors or specific anatomical structures.

Drawing biological or biomedical conclusions from ST data often requires cross-disciplinary expertise but sharing complex datasets is tedious as it often requires wrangling with customized file formats, software, and transferring of large datasets. In these cases, interactive solutions that are hostable as online resources are practical, as the data is readily visualized in a web browser, and all that has to be shared is a web link. Interactive visualization plays a key role not only for those wishing to understand the underlying biology but also for those developing these technologies and methods for analyzing data produced by such technologies. During the past years, several new methods for analyzing ST data have emerged. For example, tools for cell segmentation, identification of spatially variable genes, cell typing, data imputation, and analyses of cellcell interactions, as summarized in [19, 23]. A deeper biological understanding of the results produced by these methods can be done with the help of visual inspection. Additionally, limitations in available ground truth make methods’ quality control by visual inspection of raw data in relation to analysis results crucial.

In conclusion, the complexity and scale of ST data make interactive visualization essential, but practically difficult. We stipulate that a good tool for visualizing ST data should be able to:

**Provide multi-resolution viewing,** of gigapixel-sized images, in a manner that allows the user to flexibly view and responsively interact with the data across multiple scales. Here, a pyramidal image format is desirable as it dramatically reduces the memory requirement.

**Display large quantities (millions) of markers and regions** of various types and with adjustable size, shape, and color.

**Display features and their spatial location** in such a way that the user can simultaneously and interactively explore them in terms of spatial location and similarity.

**Be hosted,** such that interactive visualizations can conveniently be shared with the scientific community as online resources.

**Be easy to use** in terms of installation and use by a wide variety of users and systems, without expensive and costly data transfer.

### 1.1 Related work

There are several tools available for the interactive exploration of omics data, particularly with the increasing interest in ST. However, many of these tools are designed only for specific techniques. For example, STViewer [38], Spacemake [58], and 10x Genomic’s Loupe Browser [5] are all effective for exploring SST data but are not necessarily suitable for more general usage.

Viewers such as CellXGene [36], Cirrocumulus [3], and the Giotto viewer [17] are designed for the exploration of single-cell genomics and work on data where molecules have been aggregated into single-cell features. Although they provide interactive interfaces for the exploration of single-cell features in lower dimensional UMAP or t-SNE spaces, they lack functionalities for viewing individual molecular markers as well as displaying multi-layered image data.

Cytomine [32] and Visilab Viewer [30] allow the user to share large-scale interactive visualizations of gigapixel-sized images over the internet, enabling online collaboratory exploration and annotation. Their intended usage is for collaborative analysis of histopathological image data. As such, they lack functionalities for displaying markers and features, which is crucial for exploring ST data.

General purpose viewers, such as matplotlib [25], BigDataViewer [46], QuPath [9], and Napari [25] are useful for displaying markers and images but require customization to enable interactive exploration, and the image size, as well as the number of markers that can be displayed, are limited by the hardware specifications.

Vizgen’s end-to-end IST platform MERSCOPE comes with a proprietary and commercial web-viewer for interactive visualization of IST data. Using adaptive sub-sampling it can display hundreds of millions of markers on large-scale image data. However, it is strictly limited to IST data and does not allow the user to explore single-cell features in, e.g., lower-dimensional feature spaces.

The previously published TissUUmaps 1 [53] enabled interactive visualization of molecular marks on top of gigapixel-sized images. To reduce the time needed for rendering the display, markers were sub-sampled at different zoom levels. Although this improved the performance of the software, rare molecular patterns could potentially be missed. Moreover, the software was designed to be hosted on a web server, and not used directly within analysis pipelines. As such, it lacked certain usability features. For instance, it required the user to manually convert image data into a pyramidal format and configure paths to image and marker data, thereby making the software less accessible for non-programmers.

Most closely related to TissUUmaps is Vitessce [28]. Vitessce provides an extensive framework for creating and customizing interactive figures with regard to the exploration of ST data. The framework allows the user to display millions of transcripts on top of multi-scale image data, as well as display regions and single cell-feature embedding, thereby making it a powerful tool for the exploration of ST data. The framework can also be hosted on a web platform, making it easy to share interactive visualizations over the internet. However, the framework requires the user to programmatically set up the graphical user interface and configure figures by coding, making Vitessce less accessible for the non-programmer.

### 1.2 Our contribution

Summarizing the different viewers, see Table 1, we noticed that most viewers for ST are either limited to SST techniques [38, 58, 5], are not suitable for displaying millions of molecular markers [46, 36, 3, 9, 32, 52, 38], or are too general, not hostable online, and require extensive customization [25].

**Table 1:**
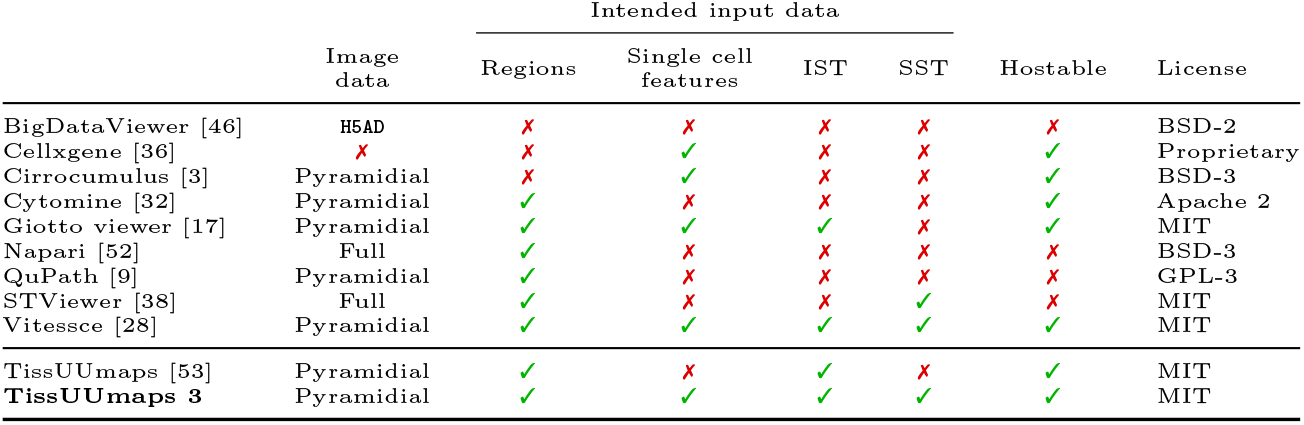
Summary of different tools for visual exploration of ST data. We noticed three approaches for loading image data: Pyramidal loading, where the same image is loaded using different resolutions depending on the zoom-level; HDF5 loading, where HDF5 file format is used for quick retrieval of large-scale data; Full loading, where the entire image is loaded into memory. The tools are designed for different types of input data: Regions, such as polygons used for delineating segmented-cells. Single cell features, such as mRNA compositions within cells, that are displayed in a lower-dimensional 2D space, and can be interactively explored; Imaging-based spatial transcriptomics (IST) data, where markers for individual transcripts/molecules are displayed in high quantities and can be interactively explored; Sequencing-based spatial transcriptomics (SST) data, where transcripts are first captured in situ and sequenced ex situ, resulting in high-dimensional data. Some tools are developed using web-based frameworks and are hostable as web-pages, allowing easy sharing of interactive visualizations as online resources. Most tools come with an open source license.

As the adoption for ST continues to grow, and now with newly emerged end-to-end systems for IST (such as Vizgen’s MERSCOPE and 10x Genomic’s Xenium), we believe that robust open-source interactive viewers for ST data are essential for researchers studying large-scale spatial omics.

In this paper, we present a web-based viewer that can handle tens of millions of unique markers, and multiple samples, without lag and at interactive frame rates, and without relying on data sub-sampling. Our contributions include maximizing the utilization of web browser capabilities, by leveraging WebGL for efficient GPU-based rendering of markers, and by optimizing how marker location and attributes are stored in memory. In addition to markers, TissUUmaps 3 can also display pie charts, graphs, and regions. Furthermore, we have structured TissUUmaps 3 to be compatible with a wide range of raw and processed data formats used in the community and built a plugin engine to provide a range of interactive analysis tools.

## 2 Methods

The improved TissUUmaps 3 implementation includes structural as well as functional changes, with the primary aim of making interactive viewing of large datasets fast and flexible. Here we describe the different implementation improvements in more detail.

### 2.1 Web based interface

The core of TissUUmaps 3 is a web-based interface developed in pure JavaScript and HTML. The JavaScript part depends on multiple open source libraries and technologies: WebGL to display markers with GPU acceleration, OpenSeadragon to display pyramidal images in Deep-Zoom format, PapaParse [24] to load and parse CSV files, h5wasm [39] to load and parse HDF5 files, D3 [16] to display regions, jquery-chosen [1] and jquery-autocomplete [2] for user-friendly inputs. The source code for TissUUmaps-core is available on the TissUUmapsCore GitHub repository at https://github.com/TissUUmaps/TissUUmapsCore. Compared to TissUUmaps 1, the main changes on the core web interface are:

#### Improved loading of CSV files

Browsers including Google Chrome and Mozilla Firefox typically limit the allocation and usage of heap memory to 4 GB per browser tab [21]. In TissUUmaps 1, the whole CSV file was loaded as a text string into memory and each row was parsed as an object, which could quickly exceed the browser memory limit for datasets with millions of points and many columns. To allow parsing of large CSV files, TissUUmaps 3 uses the JavaScript library PapaParse to stream and process data in chunks. Processed data is stored in a struct-of-arrays (SoA) layout, which allows columns with numeric data to be converted into typed arrays for lower memory consumption compared to the array-of-structs (AoS) layout used in TissUUmaps 1. Columns detected as containing only numeric values are converted and stored in Float64Array format during streaming and after loading. Using an SoA layout for marker data also has the added benefit of making TissUUmaps 3 more suitable for selectively loading columns from HDF5 files (see next item).

#### Loading HDF5 and anndata files

The anndata [60] (annotated data) file format enables flexible storing and bookkeeping of high-dimensional annotated data. As such, it has been adopted by a range of tools in the single cell and spatial omics community, reviewed in [29]. Using the h5py library on the Python side, and the h5wasm library on the JavaScript side, TissUUmaps 3 can now load anndata files, making it possible to directly and interactively explore statistical results produced by tools such as Squidpy [40] and Scanpy [64]. The anndata format is based on the Hierarchical Data Format (HDF5), enabling efficient selective loading of data columns from the HDF5 file. TissUUmaps 3 can also load HDF5 files that are not formatted as anndata files. Columns with numeric values are read directly as typed arrays which might have lower precision than 64 bits, reducing memory usage. Also, supporting HDF5 in TissUUmaps 3 gives the possibility to selectively load columns and work with very large datasets that would otherwise not fit in memory if the whole file was loaded.

#### Loading regions with GeoJSON

The file format for reading regions has been changed from a previously used in-house format [53] to the more general and widely adopted format GeoJSON [13]. Regions are displayed as polygons in a Scalable Vector Graphics (SVG) canvas.

#### WebGL rendering

TissUUmaps 1 [53] used the SVG canvas to display both regions and markers. To reduce the computational cost of rendering the markers as vector graphics, the markers were inserted into a quad-tree data structure and sub-sampled to provide a global overview of the spatial distribution. However, as mentioned above, sub-sampling may result in hiding important information from rare markers. We have therefore rewritten TissUUmaps 3 to make use of GPU acceleration through We-bGL, which allows for more efficient use of the computer resources. All markers can now be concurrently and quickly displayed, which gives the user a more accurate idea of the spatial distribution.

Two methods for marker rendering are implemented in TissUUmaps 3: The first method (point sprites) uses the native POINTS primitive of We-bGL, which allows efficient rendering because only one vertex per marker needs to be processed by the GPU during drawing. However, WebGL restricts the maximum point sprite size to 256× 256 pixels, which can be a limitation and issue when displaying large markers such as pie-charts or when viewing TissUUmaps on a high-resolution screen. The second method (instancing) uses the instancing feature of WebGL 2.0 to draw two triangles per marker; this allows arbitrarily large markers to be displayed, but at the cost of an increased number of vertices the GPU needs to process during drawing.

The vertex format used for rendering markers with WebGL requires 32 bytes of memory per marker on the GPU; this includes X and Y coordinates and marker index for selection, as well as other attributes (color, opacity, group ID, etc.) depending on the visualization parameters set by the user. In the special case of pie-chart visualization, marker data will be duplicated to display each individual pie-chart sector, which will increase GPU memory usage. To avoid having to allocate large temporary buffers in browser memory when transferring processed marker data to GPU memory, uploading to vertex buffers is performed in chunks, using the `bufferSubData()` function in WebGL. A chunk size of 100000 markers is used in the current version of TissUUmaps 3.

Finally, alpha blending is used for anti-aliasing and displaying markers with opacity.

#### Network diagram visualization

In addition to markers, TissUUmaps 3 can be used to display network diagrams (also called graphs), by adding links (edges) between markers (nodes). As for markers, the drawing of edges is implemented in WebGL and therefore benefits from GPU acceleration. Each edge is rendered by instancing two triangles, which allows for antialiasing and setting the thickness of lines. TissUUmaps 3 can thus display millions of anti-aliased edges between markers.

### 2.2 Python web server

While TissUUmaps 1 was only a static web interface, requiring a large amount of pre-processing and installation steps on the user side, TissUUmaps 3 is now embedded dynamically in a Python module to allow simplicity in installation and flexibility of use. The source code for the Python TissUUmaps module can be found on the main TissUUmaps GitHub repository: https://github.com/TissUUmaps/TissUUmaps and can be installed with the standard Python package installer pip. The TissUUmaps 3 web interface is served by a Flask server containing a dynamic OpenSlide-based Deep Zoom image converter. This allows the loading of all images recognized by the OpenSlide library (.tiff, .svs, .ndpi, etc.) without the need for pre-processing. For non-pyramidal images, an automatic converter based on VIPS [33] was added to expand the number of image formats supported by TissUUmaps (BigTIFF, OME-TIFF, etc.)

### 2.3 Native desktop application

On top of the flask server, we developed a native desktop application that can be installed and run in all major operating systems (Windows, macOS, Linux). This is implemented as a Qt web engine embedding the web interface, with the flask server running in parallel. The application code is based on PyQt6 and PyQtWebEngine, and makes it easy to use TissUUmaps 3 without access to a web server.

### 2.4 Jupyter Notebook

Jupyter notebooks are a widespread tool for prototyping algorithms, displaying data, and sharing reproducible workflows [11]. The *de facto* displaying method in notebooks is using libraries such as matplotlib [25]. We developed a Python Application Programming Interface (API) that allows interactive integration of the TissUUmaps 3 web-viewer into notebooks for quickly displaying images, regions, and markers.

### 2.5 Docker container

A docker image is also available to deploy the flask server containing TissUUmaps 3 on container orchestration systems such as Kubernetes, making it possible to share TissUUmaps projects on the web from multiple users.

### 2.6 Plugin engine

TissUUmaps 3 can be extended using plugins. Writing plugins involves programming a backend (the computational part) in Python and a front end (e.g. the web interface) in JavaScript. An API is available on the client side to create input forms for the plugins in a simple way. This allows users to customize TissUUmaps 3 and create their own tools for analysis and exploration. Examples of plugins for specific applications are showcased in Section 3.3.

## 3 Results

Here we summarize the new features of TissUUmaps 3, present results from evaluating its performance, and describe new plugins, extensions, and their applications.

### 3.1 New features

#### 3.1.1 Ways of running TissUUmaps

TissUUmaps 3 can be easily installed as a native desktop application using the standard Python package manager `pip` or a regular Windows installer. It can also be deployed on a large scale using the docker image. Finally, it can be used interactively via a Python API on Jupyter Notebooks.

#### 3.1.2 New supported data types

##### Images

combining VIPS and OpenSlide libraries, TissUUmaps 3 supports now a large number of image formats: Aperio (.svs, .tif), Hamamatsu (.ndpi, .vms, .vmu), Leica (.scn), MIRAX (.mrxs), Philips (.tiff), Sakura (.svslide), Trestle (.tif), Ventana (.bif, .tif), Generic tiled TIFF (.tif), JPEG, TIFF, PNG, WebP, FITS, Matlab, OpenEXR, PDF, SVG, HDR, PPM, CSV, GIF, Analyze, NIfTI, etc.

##### Markers

TissUUmaps 3 can load data from CSV files or anndata/HDF5 files, describing, e.g., IST or SST data, making it simple to load output from software such as StarFish [44], Squidpy [40] and Giotto [17].

##### Regions

TissUUmaps 3 can load regions in GeoJSON format. As such, TissUUmaps 3 can directly import regions exported from other software supporting GeoJSON, for example, QuPath [9].

#### 3.1.3 New marker visualization options

TissUUmaps 3 can display network diagrams using lists of neighbors per marker as edges or from anndata objects containing spatial neighbor graphs pre-computed in Squidpy [40]. Output from cell typing methods such as pciSeq [48] can be presented as pie-charts within TissUUmaps, concisely displaying local cell-type probabilities, as shown in Figure 2 c.

**Figure 1:**
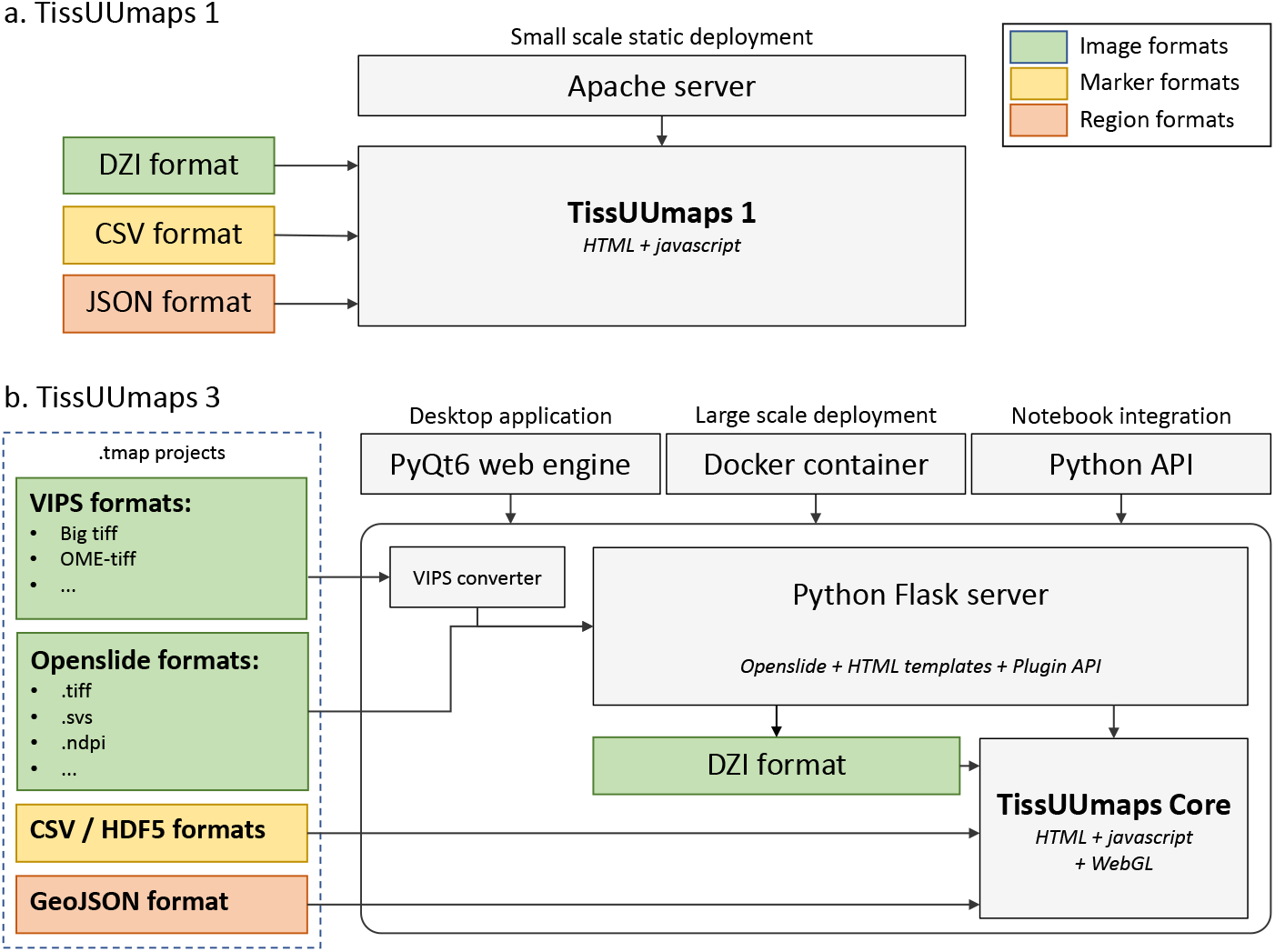
Comparison of **a.** TissUUmaps 1 infrastructure and **b.** TissUUmaps 3 infrastructure. While TissUUmaps 1 was a static web interface, TissUUmaps 3 is embedded in a Python module usable as a native desktop application, in a docker container, or interactively from a Jupyter Notebook. With its VIPS and OpenSlide image converters, TissUUmaps 3 removes the need for pre-processing images.

**Figure 2:**
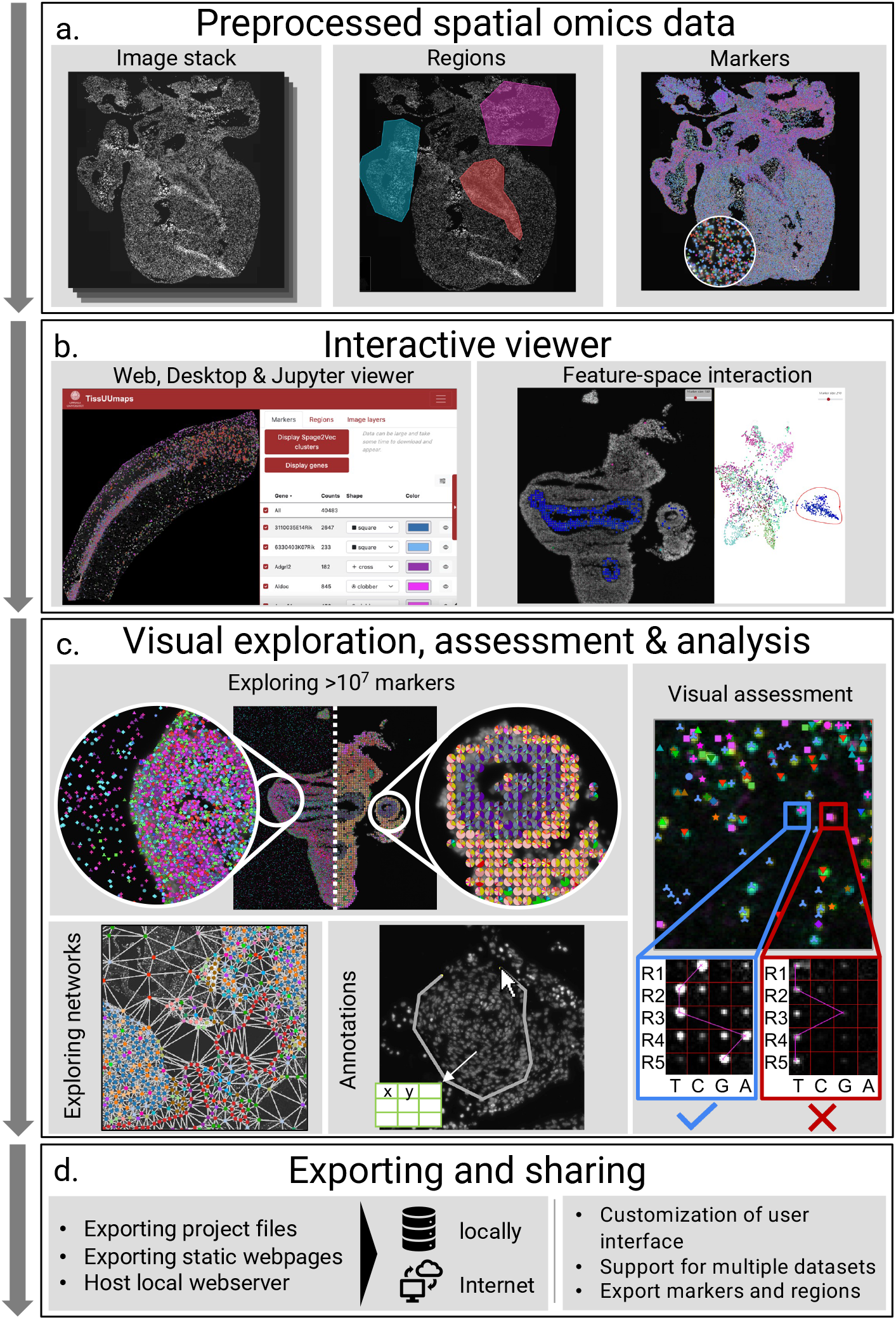
TissUUmaps is a viewer for large-scale spatial omics data. **a.** Data, in the form of regions and markers, extracted by image analysis or created by a range of different spatial statistics tools can be visualized on top of gigapixelsized image stacks. **b.** Accelerated GPU-based rendering makes image and data interaction fast, via a web or desktop viewer, or as part of a Jupyter Notebook. Interactive object selection, or ’gating’, in the feature space is directly displayed in the image space. **c.** Data can be presented and explored as markers in space or summarized in localized pie charts. Plugins for quality control enable visual assessment of raw data associated with decoded IST signals. Non-spatial data can also be interactively explored and selected, or ’gated’ signals can be overlaid on top of the original tissue sample. **d.** The data owner can export custom project files or static web pages to be run locally or over the Internet, and external users can interactively explore, manually select, and download data from regions of interest.

#### 3.1.4 User interface refinements

Intending to make the software more accessible and easier to use, we have made several refinements concerning TissUUmap’s user interface:

Image-layers can now be displayed using a composite or collection mode. In composite mode, several layers are merged using different colors into composite, multi-colored images. In collection mode, image layers are displayed side-by-side, as shown in Figure 3.
Image properties, such as opacity, saturation, brightness, contrast, color, exposure, hue, and gamma, can now be interactively adjusted from within TissUUmaps 3 (see Figure 3).
Multiple datasets (of markers) can simultaneously be loaded into TissUUmaps 3. These datasets are arranged using different tabs, as shown in Figure 3.
A configured visualization in TissUUmaps 3 can be saved as a .tmap project for later use or to share on the web.
TissUUmaps 3 comes with a capture viewport functionality, allowing the user to capture and save high-resolution screenshots of the current viewport.
The user can choose to color markers by a specified column in the loaded .csv or .hdf5 file. The column can either contain categorical labels or numeral values. In the latter case, the markers are colored using a user-selected colormap. Additional columns in the loaded file can now specify separate markers’ size, opacity, and shape.

**Figure 3:**
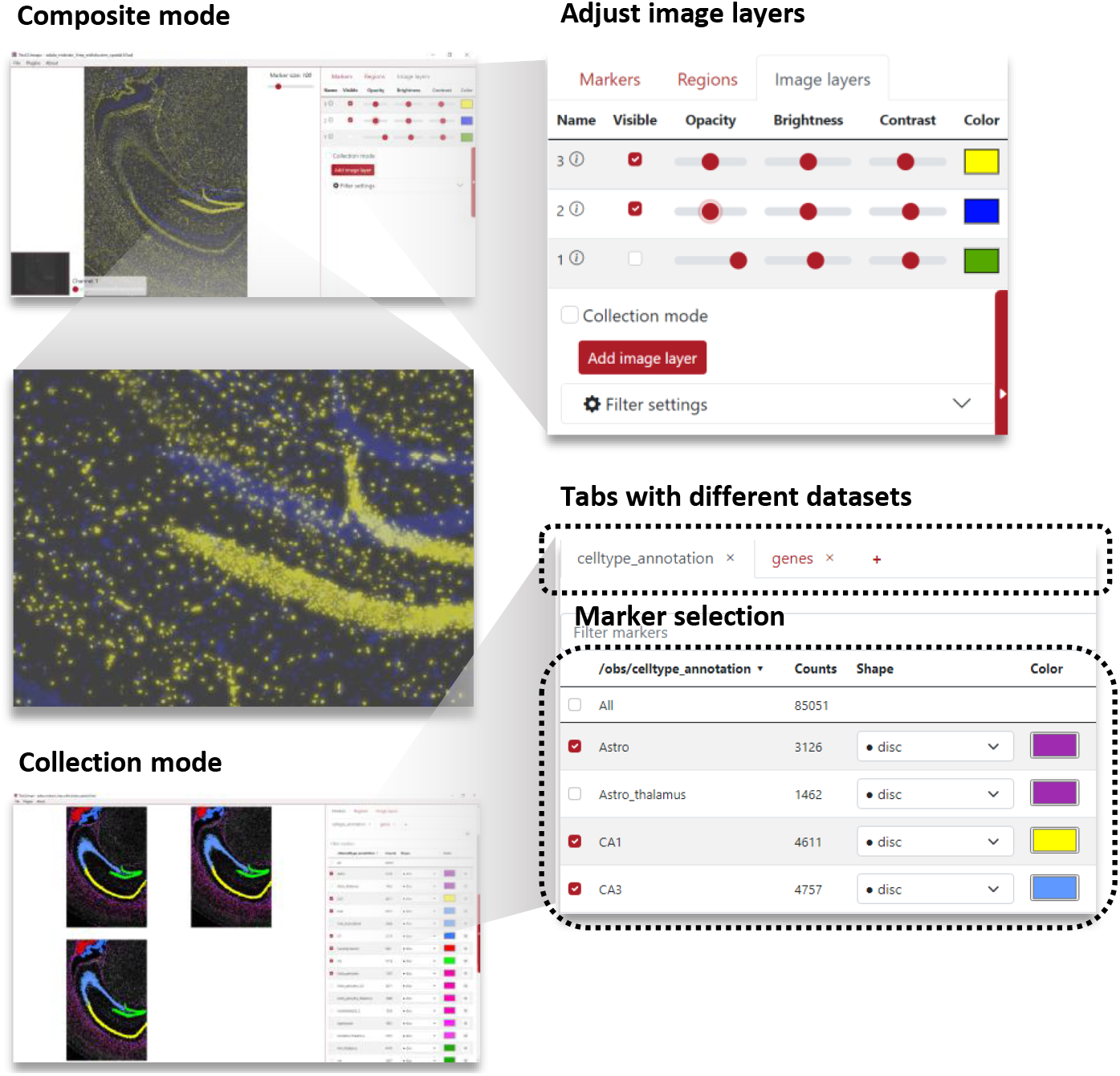
Some of the many user interface refinements done to TissUUmaps 3: Images can be displayed either in composite mode or in collection mode. In composite mode, images are blended into a composite multi-colored image. In collection mode, images are displayed side-by-side. The user can interactively adjust image properties, for instance, brightness, contrast, opacity, and color. Multiple datasets (.csv or .h5ad files) can be displayed simultaneously in TissUUmaps 3. The datasets are separated into different tabs.

### 3.2 Performance improvements

Evolving ST techniques produce huge images with millions of markers, which causes users to need increasingly more powerful computers for data exploration. For this reason, we have intensely focused on improving the performances of TissUUmaps, and here we present the results of evaluating loading time, display time, and memory usage when coping with large amounts of data.

#### 3.2.1 Loading time

First, we explored the loading time of the markers and determined how long it takes from downloading the data to displaying it. The tests were performed on a Vizgen MERFISH FFPE Human Immuno-oncology Dataset [4], and we quantified how fast the data formats used were downloaded, read, and displayed. The entire testing protocol is described in the supplementary material SI 1. Figure 4 a. shows that loading time is faster and more memory-efficient in TissUUmaps 3. Sub-sampling corresponds to using a random subset of the markers, which is quicker and decreases memory usage but displays incomplete data. HDF5 and CSV are file formats with various overheads, overall speeds, and efficiencies.

**Figure 4:**
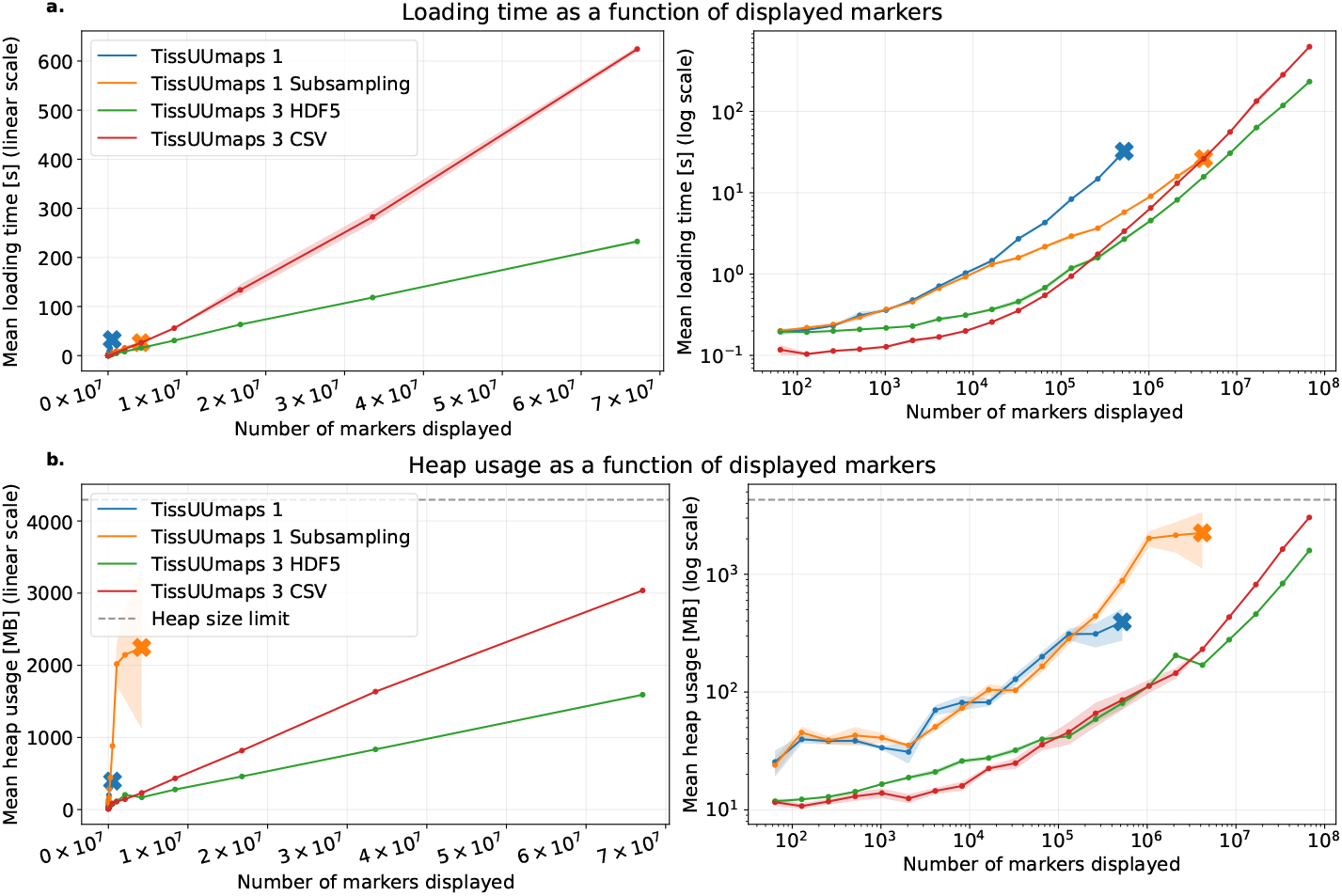
Loading time and memory usage of TissUUmaps 3 and earlier versions as a function of the number of markers displayed. The shaded area represents a 95% confidence interval around the estimated mean from 10 repeated experiments. The × symbol means that experimenting beyond that point resulted in a crash.

#### 3.2.2 Memory usage

Memory usage was performed on the same Human Immuno-oncology dataset as in the previous section. It measured the total amount of memory used by JavaScript objects as a function of displayed markers. As shown in Figure 4 b., the memory usage at run-time for both loading and visualizing markers in TissUUmaps 3 is significantly reduced compared to TissUUmaps 1 for the benchmark datasets, allowing larger datasets to be explored. The supplementary material SI 1 describes the complete testing protocol.

#### 3.2.3 Rendering time

We compare TissUUmaps 3 rendering times with its predecessor TissUUmaps 1 by measuring the average drawing time per frame for the datasets and views in Figure 5. The included datasets were three versions of the Human Immuno-Oncology dataset (with 65k, 1.0M, and 16.7M points), a Breastcancer ISS dataset [27] with 28k points (available in the TissUUmaps GitHub repository), and the Developmental Human Heart dataset [8] with 2.4M points. In addition, the quad-tree-based sub-sampling of TissUUmaps 1 was disabled in the code so that time was measured for displaying all data points. The results in Figure 5 show that the marker rendering time of TissUUmaps 3 is now much faster thanks to the GPU acceleration. This means more markers can be displayed at interactive frame rates without resorting to techniques such as sub-sampling.

**Figure 5:**
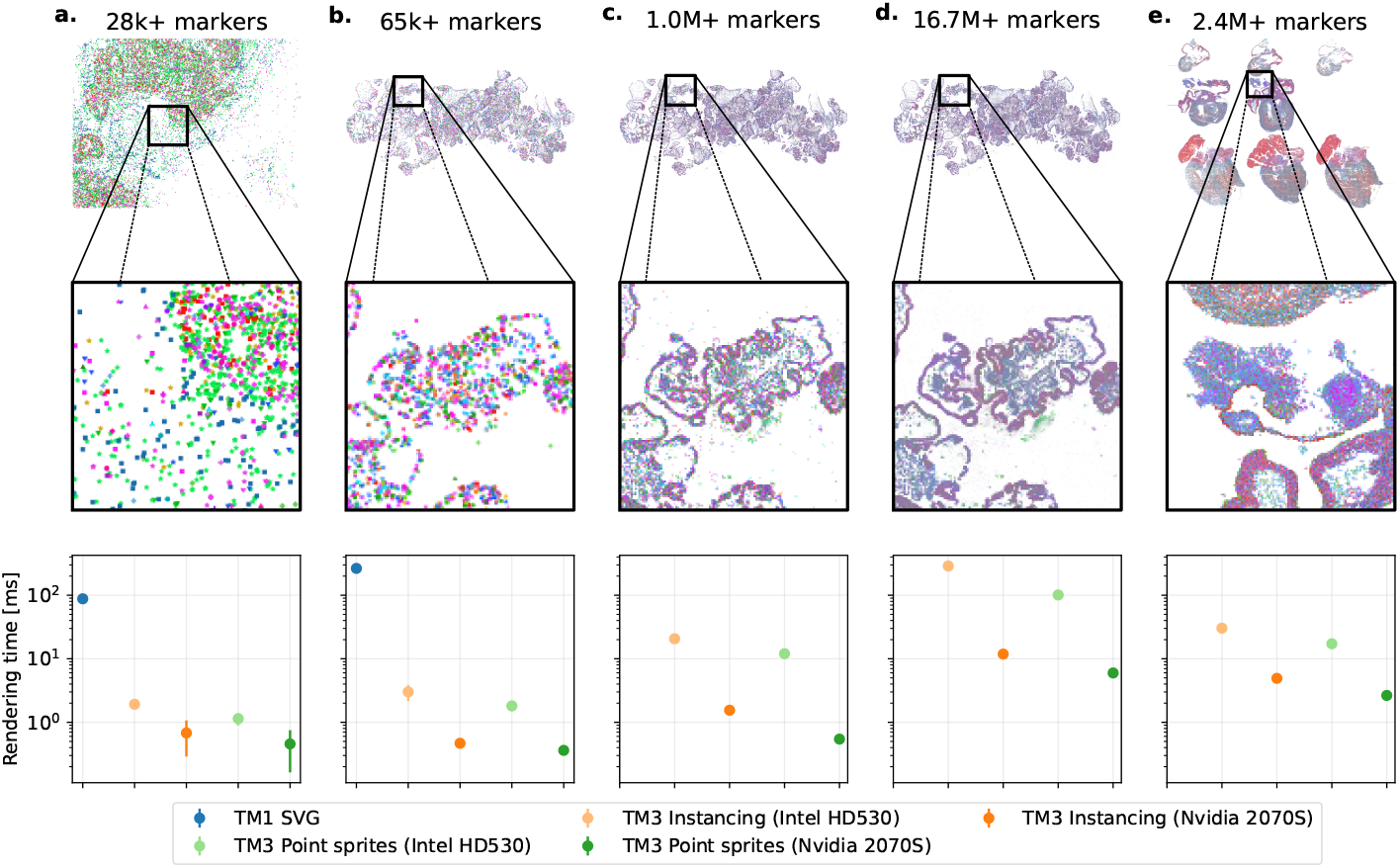
Frame times (inverse of frame-rate) over multiple datasets, GPUs, and display techniques. TissUUmaps 1 could not render the datasets c–e with interactive sub-sampling disabled. The bars represent a 95% confidence interval around the estimated means from 10 repeated experiments.

It should be noted that some browsers, including Google Chrome and Mozilla Firefox, implement WebGL on top of the software layer ANGLE, which does not support native point sprites in some cases (on Windows with the default DirectX backend) and instead emulate this feature by drawing two triangles per point [20]. We measured frame times for both rendering methods (point sprites and instancing) of TissUUmaps 3 in Google Chrome on Linux, where native point sprites are supported. To see how rendering time also scales with increased GPU performance, measurements were further repeated on a low-end integrated GPU (Intel HD530) and a higher-end discrete GPU (NVIDIA RTX 2070 Super, with 8 GB of VRAM).

Additional details about how frame times were measured for both TissUUmaps 3 and TissUUmaps 1 can be found in the supplementary material SI 1.

### 3.3 Plugins, extensions, and their applications

As mentioned in Section 2.6, TissUUmaps 3 can now be extended with plugins that typically consist of a backend computational part written in Python and a front-end built on the TissUUmaps 3 web interface in JavaScript. Therefore, we provide a plugin template, including an API and several examples of plugins, all collected at https://tissuumaps.github.io/TissUUmaps/plugins/. This allows users to package their code and customize an accompanying parameter selection interface, view-port, and interface for user interaction, as illustrated in Figure 6 a. Examples of provided plugins are described below.

**Figure 6:**
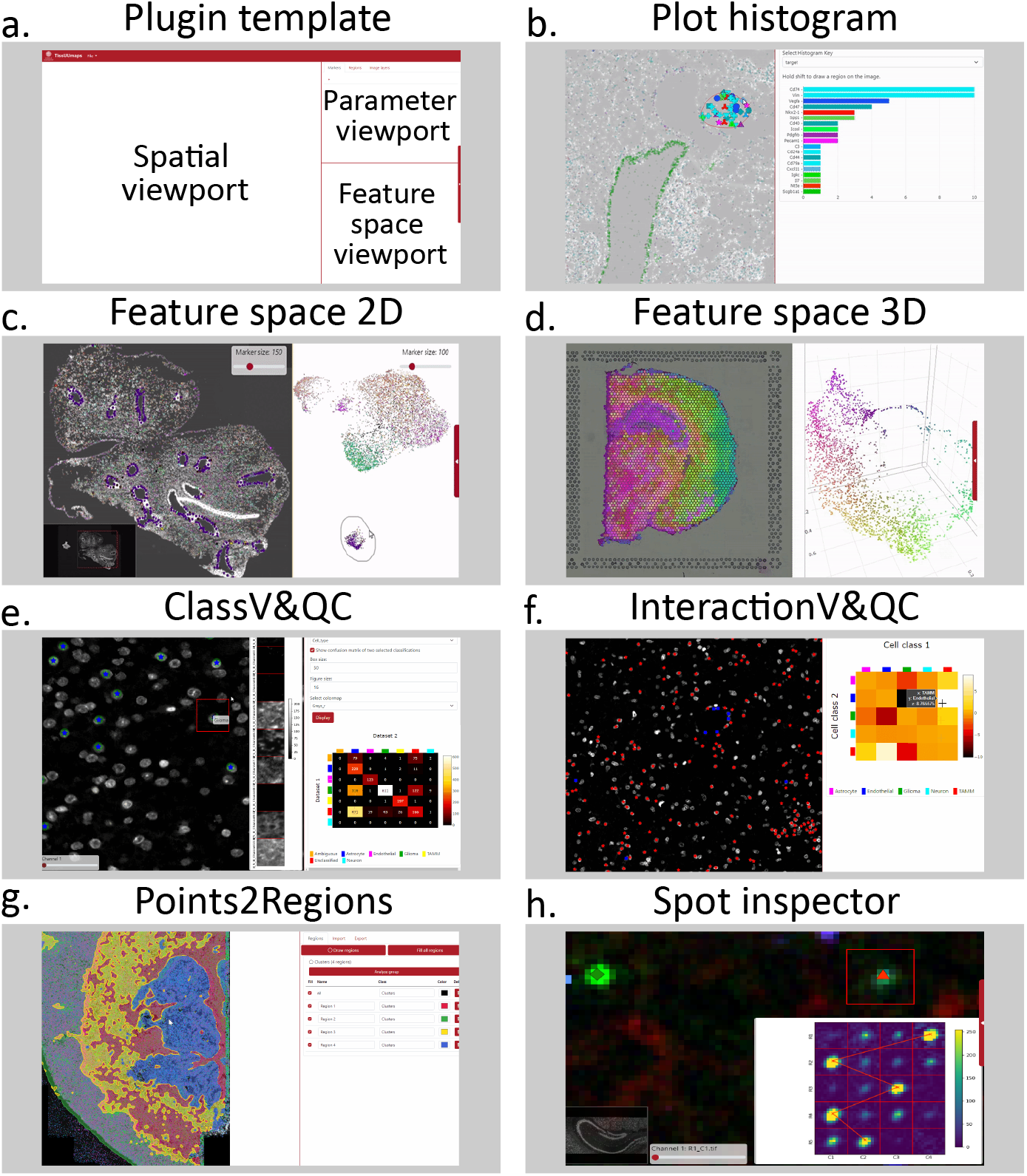
TissUUmaps 3 plugins; Users may create their own plugins using a template (**a**). A range of plugins for specific tasks are provided at https://tissuumaps.github.io/TissUUmaps/plugins/ (**b-h**).

#### 3.3.1 Plugin for an interactive view of point distributions

The `PlotHistogram` plugin provides a quick overview of ST data distribution, in the form of a histogram, within a manually drawn region of interest. This makes it easy to visually explore similarities and differences within different parts of a tissue section, selecting a region in the viewport, as shown in Figure 6 b, left, and the corresponding histogram of counts per gene will show up to the right.

#### 3.3.2 Plugin for 2D feature space exploration

A spatial omics data point is typically associated with two different attributes: a spatial position and a feature descriptor. The type of feature descriptor varies between different techniques. For example, in an IST experiment, the descriptor is usually a type of compositional vector, describing the composition of mRNA molecules in a small neighborhood, whereas in a multiplexed immunohistochemistry experiment, the descriptor may be a vector describing the fluorescent signal in different image channels, within e.g. a single cell. Downstream analysis of such descriptors usually leads to the identification of various entities in the tissue, for example, the identification of different cell types. To construct and validate biological hypotheses, it is important to explore data both in terms of physical location and feature similarity. Therefore, we have developed a plugin called `FeatureSpace`.

The plugin starts with the user computing a 2D projection of the feature descriptor using standard dimensionality reduction techniques such as UMAP [10] or t-SNE [59]. `FeatureSpace` allows the user to simultaneously display the physical coordinates of each data point on top of image data together with the 2D projection of the feature descriptors. The user can then interactively select regions of interest in either physical space or in feature space, see Figure 6 c. The plugin will automatically highlight where the selected data points are located in both of the spaces, enabling efficient exploratory analysis of spatial omics data.

#### 3.3.3 Plugin for 3D feature space visualization

A 2D view of a multi-dimensional feature space is often limiting, hiding subclusters. Therefore, we have developed a plugin called `FeatureSpace3D`, which makes it possible to view and rotate a 3D representation of a feature space. Figure 6 d illustrates a 3D feature space color-coded according to RGB color space, together with a 2D representation of the corresponding spatial coordinates on top of a tissue section, as described in [15].

#### 3.3.4 Classification Visualization and Quality Control

Classification is a fundamental step when working with spatial omics data for, e.g., understanding the distribution of cell types in a tissue. Classification techniques and/or parameter settings may result in varying classification results. We created the Classification Visualization and Quality Control, or `ClassV&QC` plugin to visualize and set side-by-side results of two techniques for classification, as described in more detail in [12]. Briefly, when comparing the output of two classification techniques, one output is displayed as large circles on the spatial viewport, while the output of the second technique is displayed as small stars on top of the circles. This allows the user to see both classification results at the same time. If the two methods have the same marker coordinates in the .csv files, the plugin can display an interactive confusion matrix where the user can click on the elements of the matrix, and only markers counted in that matrix element are displayed on the Spatial viewport. With this, the user can inspect the classification or the misclassification of two approaches from the spatial point of view. Additionally, the built-in `Signal Inspector` plugin makes it possible to investigate the result in relation to cut-outs of the local staining patterns, to better evaluate which classification technique is more correct, as shown in Figure 6 e.

#### 3.3.5 Interaction Visualization and Quality Control

The spatial proximity of two different categories of markers can be quantified by a neighborhood enrichment test. This test assigns higher positive values to a pair of marker categories that are spatially closer to each other than they would be by random, values around zero to a pair of marker categories that are randomly distributed in relation to one another, and lower negative values to a pair of marker categories that are spatially further away from each other than they would be by random. The results of the neighborhood enrichment test can be exported as a matrix where columns and rows represent different marker categories, as described in more detail in [12]. We created an Interaction Visualization and Quality Control, or `InteractionV&QC` plugin, which can load a neighborhood enrichment matrix and make it interactive, so the user can click on the elements of the matrix and only those two corresponding marker categories are displayed on the Spatial viewport. Through this, the user can investigate quantified proximity from the spatial point of view, as illustrated in Figure 6 f.

#### 3.3.6 Plugin for interactive quality assessment for multiplexed image data

Assessing the quality of fluorescent multiplexed image data is a tedious task as it requires the inspection of signals from multiple image layers (such as different fluorescent channels collected from various staining rounds). Therefore, we have developed a plugin named `Signal Inspector` that allows the user to interactively click anywhere on an image being shown in TissUUmaps 3. The plugin will automatically open a subfigure displaying image content from all remaining image layers. We believe this plugin to be particularly useful for inspecting and assessing the quality of the combinatorial labeling schemes used in IST techniques. As such, the plugin can also display results from decoding methods as traces connecting fluorescent signals across various image layers, see Figure 6 h.

#### 3.3.7 Plugin for unsupervised data clustering

A single IST experiment can target hundreds of different types of mRNA species (genes) and generate several million molecular markers. Molecular patterns, among this large quantity of markers, become overwhelming if all markers are displayed at once – there is simply too much information at hand to visually interpret the data. A common strategy to dissect the data is to reduce the number of dimensions, for instance, via clustering. We have therefore developed a plugin to TissUUmaps 3, named `Points2Regions`, that allows the user to interactively cluster the molecular markers into distinct regions with similar molecular compositions [7], as shown in Figure 6 g. Since clustering and visualization have been optimized for speed, the resulting visualization of clusters in response to parameter settings takes a few seconds for reasonably sized datasets, making parameter exploration easy.

### 3.4 Interfacing with other software

There are many powerful tools for the analysis of microscopy data, producing regions and markers as output. We provide examples for some widely used open source tools and show how they can interface with visual exploration via TissUUmaps 3:

QuPath [9] is an open-source tool for digital pathology, supporting the classification, annotation, and segmentation of different cells and tissue regions. Such results can be visualized as markers or regions in TissUUmaps by exporting the QuPath results to a GeoJSON file which in turn can be read directly into TissUUmaps.

Napari [52] is a python-based multi-dimensional viewer, with a large number of associated packages, e.g., for spatial statistics via Squidpy [40], and cell segmentation via Cellpose [57]. Output from Napari projects, including images, annotations, or labeling (as masks and vector graphics), as well as point clouds, can be loaded in TissUUmaps 3 via a Napari-TissUUmaps 3 plugin available on the Napari Hub, as described in https://tissuumaps.github.io/tutorials/#napari.

CellProfiler [14, 35] is a popular tool for performing high-throughput analyses of microscopy image data. CellProfiler usually outputs results in the form of CSV files, containing locations, and associated feature measurements, that can be loaded as markers directly in TissUUmaps 3, and viewed by selecting columns representing coordinates and features of interest (including marker type and color) in the TissUUmaps 3 GUI.

Fiji [51], another popular tool for image analysis, bundles a lot of plugins that facilitate scientific image processing, including image registration. An example of how the registration functionality of Fiji can be output to fit TissUUmaps 3 is described at https://github.com/TissUUmaps/TissUUmaps/blob/master/examples/fiji_alignment_dot_extraction.py.

## 4 Discussion and Conclusion

Spatial transcriptomics was featured as the method of the year in Nature 2020 [34], and a range of ST techniques are now made broadly available by commercial companies. At the same time, the development of new and improved techniques results in output data quickly growing richer in terms of the number of transcripts per unit area, the number of different transcripts detected in parallel, and the dimensions of analyzed tissue samples, calling for tools that can help us comprehend these enormous datasets. A first step is often visualization, and here we believe the efficient rendering and interactive data exploration provided by TissUUmaps 3 is crucial. As the adoption of ST methods continues to grow there may be several areas where TissUUmaps 3 could be expanded with new functionality. Combined analysis of ST data and tissue morphology, building on developments in AI techniques for digital pathology, stands out as a potential area of improvement. The large file sizes associated with tissue specimens pose a challenge for image analysis, as system memory typically is not sufficient to load the entire image at once. This means that tools producing measurements to be used as input for TissUUmaps 3 may pose a bottleneck that could be avoided by adopting packages such as Dask [49], as a means of handling such images by loading subsections of an image on-demand.

We are also following the development of the Open Microscopy Environment - Next-Generation File Formats (OME-NGFF) for cloud-based data storage [37], and we are considering expanding TissUUmaps 3 to directly load such data, in line with the OME-NGFF idea that global, collaborative discovery as opposed to the post-publication, ”data-on-request” mode of operation is the path forward. This includes considering cloud storage systems such as Google Cloud Storage (GCS) and Amazon Simple Storage Service (Amazon S3).

So far, we have primarily focused on optimizing TissUUmaps 3 as a data viewer. However, with our recent development of the Points2Regions plugin [7], it has become clear that interactive tools that also include a speed-optimized computational step are very valuable for data exploration. With Points2Regions, the user can vary parameters such as the radius of the local neighborhood, and directly explore ST composition ranging from single cells to larger functional tissue regions. We foresee that the integration of TissUUmaps 3 with Jupyter Notebooks may be a good environment for developing more such tools that could eventually be integrated as plugins.

Many different questions can be posed to the same dataset, and data sharing according to the FAIR principles [63], aiming to improve the Findability, Accessibility, Interoperability, and Reuse of digital assets, is highly relevant for ST data. Visualizing data prior to download is important for judging the relevance of a dataset, and future development for TissUUmaps 3 is a simplified targeted download of, e.g., regions and markers of interest, reducing the time and cost of data sharing. We are already using TissUUmaps 3 as a means of sharing published datasets, as exemplified in our gallery at https://tissuumaps.github.io/gallery/, providing IST and ST data from the Developmental Human Cell Atlas lung [54] and heart [43, 8].

In this manuscript, we presented an updated version of TissUUmaps that has been optimized to ensure that it can provide suitable usage performance even as ST datasets are growing exponentially, accommodating the continuous development of ST techniques. We demonstrated that TissUUmaps 3 provides significantly improved performance for large multiplex datasets as compared to previous versions, at the same time as we have increased ease of use and options for customization. We envision TissUUmaps 3 to be a valuable resource for researchers studying large-scale spatial omics data, and we believe that it will contribute to the broader dissemination and flexible sharing of this type of data.

## 5 Funding Received

This work was supported by the European Research Council via ERC Consolidator (grant number CoG 682810), the Swedish Foundation for Strategic Research (grant numbers SB160046 and BD150008), and the SciLifeLab BioImage Informatics Facility / NBIS.

## 7 Acknowledgements

We would like to thank all our collaborators for their valuable input during the development of TissUUmaps.

## Supplementary Information

### SI 1 Benchmarking protocols and specifications

#### Loading time and memory usage

To measure loading time in the benchmarks in Section 3.2, the following hardware and software specifications were used:

- A simulated, fixed network bandwidth of 100Mbit/s, using a custom Network Throttling Profile of the Chrome developer tools.
- An HTTP server running locally with support for partial requests from the client to remove internet connection variability and allow range access for HDF5 datasets.
- Google Chrome browser v108.0.5359.100 running on Windows 10 64-bit, with an x64-based processor in-client requests, 1GHz, integrated GPU (Intel UHD Graphics 620), and 32 GB of RAM.

Each measure was performed ten times. Time was measured using the JavaScript `time` function from the start of data downloading until the markers appeared in the viewport. Memory was measured using the JavaScript function `performance.memory.usedJSHeapSize` in Chrome. The browser tab was closed and reopened between runs to get the most reliable measure of the used heap size. It can be noted that this does not avoid all variations between runs, as the memory usage in JavaScript depends on multiple factors.

The datasets used were sub-sampled randomly from *Vizgen MERFISH FFPE Human Immuno-oncology Data Set, May 2022: Human Ovarian Cancer Patient 2, Slice 1*, with all successive powers of two from 2^6^ to 2^36^.

#### Rendering time

To measure rendering time in the benchmarks in Section 3.2, the following hardware and software specifications were used:

- A laptop with an Intel Core i7-6700HQ CPU, integrated GPU (Intel HD Graphics 530), and 32 GB of RAM.
- A desktop with an Intel Core i5-6600K CPU, discrete GPU (NVIDIA RTX 2070 Super, with 8 GB of VRAM), and 32 GB of RAM.
- Google Chrome browser v108.0.5359.98 running on Ubuntu 20.04 64-bit on both computers.

For the WebGL-based marker rendering in TissUUmaps 3, we used timer queries from the API extension `EXT_disjoint_timer_query_webgl2` to log GPU frame timings for the rasterization of the markers. For the SVG-based marker rendering in TissUUmaps 1, we could not directly measure frame timings in the browser. Instead, frame times were estimated from frame rates captured at different time points via the “Show frames per second (FPS) meter” overlay in the Chrome developer tools.

Average frame time over ten frames was measured for each combination of a dataset (Figure 5) and TissUUmaps version; we also repeated the measurements on both test systems for each rendering method (point sprites or instancing) of TissUUmaps 3. On the test system with the NVIDIA RTX 2070 Super GPU, the performance mode in the NVIDIA X Server Settings control panel was set to “Prefer Maximum Performance” for more consistent frame timings for the smaller datasets. In Figure 5, confidence intervals for the average frame times are also presented.

## References

[1] GitHub - harvesthq/chosen: Deprecated - Chosen is a library for making long, unwieldy select boxes more friendly. — github.com. https://github.com/harvesthq/chosen. [Accessed 13-Dec-2022].

[2] GitHub - jquery/jqueryui.com: jQuery UI web site content — github.com. https://github.com/jquery/jqueryui.com. [Accessed 14-Dec-2022].

[3] aGitHub - lilab-bcb/cirrocumulus: Bring your single-cell data to life — github.com. https://github.com/lilab-bcb/cirrocumulus. [Accessed 30-Nov-2022].

[4] Vizgen MERFISH FFPE Human Immuno-oncology Data Set, may 2022: Human ovarian cancer patient 2, slice 1.https://info.vizgen.com/ffpe-showcase?submissionGuid=689a82e4-f76a-42da-a0c6-b95a987a7adc. [Accessed 21-Dec-2022].

[5] 10x Genomics. Loupe Browser. https://www.10xgenomics.com/products/loupe-browser. [Accessed 30-Nov-2022].

[6] A. C. Anderson, I. Yanai, L. R. Yates, L. Wang, A. Swarbrick, P. Sorger, S. Santagata, W. H. Fridman, Q. Gao, L. Jerby, B. Izar, L. Shang, and X. Zhou. Spatial transcriptomics. Cancer Cell, 40(9):895–900, 2022.

[7] A. Andersson, A. Behanova, C. Avenel, C. Wählby, and F. Malmberg. Points2regions: Fast interactive clustering of in situ transcriptomics data. bioRxiv, 2022.

[8] M. Asp, S. Giacomello, L. Larsson, C. Wu, D. Fürth, X. Qian, E. Wärdell, J. Custodio, J. Reimegård, F. Salmén, C. Österholm, P. L. Ståhl, E. Sund-ström, E. Åkesson, O. Bergmann, M. Bienko, A. Månsson-Broberg, M. Nilsson, C. Sylvén, and J. Lundeberg. A spatiotemporal organ-wide gene expression and cell atlas of the developing human heart. Cell, 179(7):1647–1660.e19, Dec. 2019.

[9] P. Bankhead, M. B. Loughrey, J. A. Fernández, Y. Dombrowski, D. G. McArt, P. D. Dunne, S. McQuaid, R. T. Gray, L. J. Murray, H. G. Coleman, et al. Qupath: Open source software for digital pathology image analysis. Scientific reports, 7(1):1–7, 2017.

[10] E. Becht, L. McInnes, J. Healy, C.-A. Dutertre, I. W. Kwok, L. G. Ng, F. Ginhoux, and E. W. Newell. Dimensionality reduction for visualizing single-cell data using umap. Nature biotechnology, 37(1):38–44, 2019.

[11] M. Beg, J. Taka, T. Kluyver, A. Konovalov, M. Ragan-Kelley, N. M. Thiery, and H. Fangohr. Using jupyter for reproducible scientific workflows. Computing in Science and Engineering, 23(2):36–46, 2021.

[12] A. Behanova, C. Avenel, A. Andersson, E. Chelebian, A. Klemm, L. Wik, A. Ostman, and C. Wählby. Tissuumaps 3 tools for visualization & quality control in large-scale multiplex tissue analysis. bioRxiv, 2022.

[13] H. Butler, M. Daly, A. Doyle, S. Gillies, S. Hagen, and T. Schaub. The geojson format. Technical report, 2016.

[14] A. E. Carpenter, T. R. Jones, M. R. Lamprecht, C. Clarke, I. H. Kang, O. Friman, D. A. Guertin, J. H. Chang, R. A. Lindquist, J. Moffat, et al. Cellprofiler: image analysis software for identifying and quantifying cell phenotypes. Genome biology, 7(10):1–11, 2006.

[15] E. Chelebian, C. Avenel, K. Kartasalo, M. Marklund, A. Tanoglidi, T. Mirtti, R. Colling, A. Erickson, A. D. Lamb, J. Lundeberg, et al. Morphological features extracted by ai associated with spatial transcriptomics in prostate cancer. Cancers, 13(19):4837, 2021.

[16] D.-D. Documents. GitHub - d3/d3: Bring data to life with SVG, Canvas and HTML. — github.com. https://github.com/d3/d3. [Accessed 13-Dec-2022].

[17] R. Dries, Q. Zhu, R. Dong, C.-H. L. Eng, H. Li, K. Liu, Y. Fu, T. Zhao, A. Sarkar, F. Bao, et al. Giotto: a toolbox for integrative analysis and visualization of spatial expression data. Genome biology, 22(1):1–31, 2021.

[18] C.-H. L. Eng, M. Lawson, Q. Zhu, R. Dries, N. Koulena, Y. Takei, J. Yun, C. Cronin, C. Karp, G.-C. Yuan, et al. Transcriptome-scale super-resolved imaging in tissues by rna seqfish+. Nature, 568(7751):235–239, 2019.

[19] S. Fang, B. Chen, Y. Zhang, H. Sun, L. Liu, S. Liu, Y. Li, and X. Xu. Computational approaches and challenges in spatial transcriptomics. Genomics, Proteomics & Bioinformatics, 2022.

[20] Google. GitHub - ANGLE: Almost Native Graphics Layer Engine — github.com. https://github.com/google/angle/blob/66ee308c4cd3aa7ea4d66219c81a7e6680cfcc0c/src/libANGLE/renderer/d3d/d3d11/Renderer11.cpp#L1853. [Accessed 13-Dec-2022].

[21] Google. GitHub - Chromium: Tab memory size limit — github.com. https://github.com/chromium/chromium/blob/7073a66a6f227ed904b53c353d1340e4a322f3f2/sandbox/constants.h#L20. [Accessed 21-Dec-2022].

[22] D. Gyllborg, C. M. Langseth, X. Qian, E. Choi, S. M. Salas, M. M. Hilscher, E. S. Lein, and M. Nilsson. Hybridization-based in situ sequencing (hybiss) for spatially resolved transcriptomics in human and mouse brain tissue. Nucleic acids research, 48(19):e112–e112, 2020.

[23] A. A. Heydari and S. S. Sindi. Deep learning in spatial transcriptomics: Learning from the next next-generation sequencing. Mar. 2022.

[24] M. Holt. Papaparse. https://github.com/mholt/PapaParse. [Accessed 30-Nov-2022].

[25] J. D. Hunter. Matplotlib: A 2d graphics environment. Computing in science & engineering, 9(03):90–95, 2007.

[26] N. Ji and A. Van Oudenaarden. Single molecule fluorescent in situ hybridization (smfish) of c. elegans worms and embryos. 2012.

[27] R. Ke, M. Mignardi, A. Pacureanu, J. Svedlund, J. Botling, C. Wählby, and M. Nilsson. In situ sequencing for rna analysis in preserved tissue and cells. Nature methods, 10(9):857–860, 2013.

[28] M. S. Keller, I. Gold, C. McCallum, T. Manz, P. V. Kharchenko, and N. Gehlenborg. Vitessce: a framework for integrative visualization of multimodal and spatially-resolved single-cell data. OSF Preprints, Oct. 2021.

[29] I. Kleino, P. Frolovaitė, T. Suomi, and L. L. Elo. Computational solutions for spatial transcriptomics. Computational and Structural Biotechnology Journal, 20:4870–4884, 2022.

[30] N. Lajara, J. L. Espinosa-Aranda, O. Deniz, and G. Bueno. Optimum web viewer application for dicom whole slide image visualization in anatomical pathology. Computer Methods and Programs in Biomedicine, 179:104983, 2019.

[31] J. H. Lee, E. R. Daugharthy, J. Scheiman, R. Kalhor, T. C. Ferrante, R. Terry, B. M. Turczyk, J. L. Yang, H. S. Lee, J. Aach, et al. Fluorescent in situ sequencing (fisseq) of rna for gene expression profiling in intact cells and tissues. Nature protocols, 10(3):442–458, 2015.

[32] R. Marée, L. Rollus, B. Stévens, R. Hoyoux, G. Louppe, R. Vandaele, J.-M. Begon, P. Kainz, P. Geurts, and L. Wehenkel. Collaborative analysis of multi-gigapixel imaging data using cytomine. Bioinformatics, 32(9):1395–1401, Jan. 2016.

[33] K. Martinez and J. Cupitt. Vips-a highly tuned image processing software architecture. In IEEE International Conference on Image Processing 2005,volume 2, pages II–574. IEEE, 2005.

[34] V. Marx. Method of the year: spatially resolved transcriptomics. Nature Methods, 18(1):9–14, Jan. 2021.

[35] C. McQuin, A. Goodman, V. Chernyshev, L. Kamentsky, B. A. Cimini, K. W. Karhohs, M. Doan, L. Ding, S. M. Rafelski, D. Thirstrup, et al. Cellprofiler 3.0: Next-generation image processing for biology. PLoS biology, 16(7):e2005970, 2018.

[36] C. Megill, B. Martin, C. Weaver, S. Bell, S. Badajoz, M. Weiden, J. Kiggins, J. Freeman, Fionagriffin, Bmccandless, M. Kinsella, Snyk Bot, Prete, A. Philipp, M. V. Muhlen, J. Taylor, I. Virshup, Gökçen Eraslan, GenevieveHaliburton, and A. Wolf. chanzuckerberg/cellxgene: Release 0.15.0, 2020.

[37] J. Moore. ome/ngff: Next-generation file format (ngff) specifications for storing bioimaging data in the cloud., 2020.

[38] J. F. Navarro, J. Lundeberg, and P. L. Ståhl. ST viewer: a tool for analysis and visualization of spatial transcriptomics datasets. Bioinformatics, 35(6):1058–1060, Aug. 2018.

[39] N. I. of Standards and Technology. GitHub - usnistgov/h5wasm: A WebAssembly HDF5 reader/writer library — github.com. https://github.com/usnistgov/h5wasm. [Accessed 13-Dec-2022].

[40] G. Palla, H. Spitzer, M. Klein, D. Fischer, A. C. Schaar, L. B. Kuemmerle, S. Rybakov, I. L. Ibarra, O. Holmberg, I. Virshup, et al. Squidpy: a scalable framework for spatial omics analysis. Nature methods, 19(2):171–178, 2022.

[41] J. Park, W. Choi, S. Tiesmeyer, B. Long, L. E. Borm, E. Garren, T. N. Nguyen, B. Tasic, S. Codeluppi, T. Graf, et al. Cell segmentation-free inference of cell types from in situ transcriptomics data. Nature communications, 12(1):1–13, 2021.

[42] G. Partel, M. M. Hilscher, G. Milli, L. Solorzano, A. H. Klemm, M. Nilsson, and C. Wählby. Automated identification of the mouse brain’s spatial compartments from in situ sequencing data. BMC biology, 18(1):1–14, 2020.

[43] G. Partel and C. Wählby. Spage2vec: Unsupervised representation of localized spatial gene expression signatures. The FEBS Journal, 288(6):1859–1870, Oct. 2020.

[44] J. M. Perkel. Starfish enterprise: finding rna patterns in single cells. Nature, 572(7770):549–549, 2019.

[45] V. Petukhov, R. J. Xu, R. A. Soldatov, P. Cadinu, K. Khodosevich, J. R. Moffitt, and P. V. Kharchenko. Cell segmentation in imaging-based spatial transcriptomics. Nature Biotechnology, 40(3):345–354, 2022.

[46] T. Pietzsch, S. Saalfeld, S. Preibisch, and P. Tomancak. Bigdataviewer: visualization and processing for large image data sets. Nature methods, 12(6):481–483, 2015.

[47] A. J. Piñeiro, A. E. Houser, and A. L. Ji. Research techniques made simple: Spatial transcriptomics. Journal of Investigative Dermatology, 142(4):993–1001.e1, 2022.

[48] X. Qian, K. D. Harris, T. Hauling, D. Nicoloutsopoulos, A. B. Munoz-Manchado, N. Skene, J. Hjerling-Leffler, and M. Nilsson. Probabilistic cell typing enables fine mapping of closely related cell types in situ. Nature methods, 17(1):101–106, 2020.

[49] M. Rocklin. Dask: Parallel computation with blocked algorithms and task scheduling. In Proceedings of the 14th python in science conference, number 130–136. Citeseer, 2015.

[50] S. G. Rodriques, R. R. Stickels, A. Goeva, C. A. Martin, E. Murray, C. R. Vanderburg, J. Welch, L. M. Chen, F. Chen, and E. Z. Macosko. Slide-seq: A scalable technology for measuring genome-wide expression at high spatial resolution. Science, 363(6434):1463–1467, 2019.

[51] J. Schindelin, I. Arganda-Carreras, E. Frise, V. Kaynig, M. Longair, T. Pietzsch, S. Preibisch, C. Rueden, S. Saalfeld, B. Schmid, et al. Fiji: an open-source platform for biological-image analysis. Nature methods, 9(7):676–682, 2012.

[52] N. Sofroniew, T. Lambert, K. Evans, J. Nunez-Iglesias, G. Bokota, G. P. na Castellanos, P. Winston, K. Yamauchi, M. Bussonnier, D. D. Pop, Z. Liu, ACS, Pam, alisterburt, G. Buckley, A. Sweet, L. Gaifas, J. Rodríguez-Guerra, L. Migas, V. Hilsenstein, J. ao Bragantini, G. R. Lee, Hector, J. Freeman, P. Boone, A. R. Lowe, C. Gohlke, L. Royer, A. PIERRÉ, and H. Har-Gil. napari/napari: 0.4.12rc2, Oct. 2021.

[53] L. Solorzano, G. Partel, and C. Wählby. TissUUmaps: interactive visualization of large-scale spatial gene expression and tissue morphology data. Bioinformatics, 36(15):4363–4365, 05 2020.

[54] A. Sountoulidis, S. M. Salas, E. Braun, C. Avenel, J. Bergenstråhle, M. Vicari, P. Czarnewski, J. Theelke, A. Liontos, X. Abalo, ž. Andrusivová, M. Asp, X. Li, L. Hu, S. Sariyar, A. M. Casals, B. Ayoglu, A. Firsova, J. Michaëlsson, E. Lundberg, C. Wählby, E. Sundström, S. Linnarsson, J. Lundeberg, M. Nilsson, and C. Samakovlis. Developmental origins of cell heterogeneity in the human lung. bioRxiv, 2022.

[55] P. L. Ståhl, F. Salmén, S. Vickovic, A. Lundmark, J. F. Navarro, J. Magnusson, S. Giacomello, M. Asp, J. O. Westholm, M. Huss, et al. Visualization and analysis of gene expression in tissue sections by spatial transcriptomics. Science, 353(6294):78–82, 2016.

[56] R. R. Stickels, E. Murray, P. Kumar, J. Li, J. L. Marshall, D. J. Di Bella, P. Arlotta, E. Z. Macosko, and F. Chen. Highly sensitive spatial transcriptomics at near-cellular resolution with slide-seqv2. Nature biotechnology, 39(3):313–319, 2021.

[57] C. Stringer, T. Wang, M. Michaelos, and M. Pachitariu. Cellpose: a generalist algorithm for cellular segmentation. Nature Methods, 18(1):100–106, 2021.

[58] T. R. Sztanka-Toth, M. Jens, N. Karaiskos, and N. Rajewsky. Spacemake: processing and analysis of large-scale spatial transcriptomics data. Giga-Science, 11, 2022.

[59] L. Van der Maaten and G. Hinton. Visualizing data using t-sne. Journal of machine learning research, 9(11), 2008.

[60] I. Virshup, S. Rybakov, F. J. Theis, P. Angerer, and F. A. Wolf. anndata: Annotated data. bioRxiv, 2021.

[61] G. Wang, J. R. Moffitt, and X. Zhuang. Multiplexed imaging of high-density libraries of rnas with merfish and expansion microscopy. Scientific reports, 8(1):1–13, 2018.

[62] X. Wang, W. E. Allen, M. A. Wright, E. L. Sylwestrak, N. Samusik, S. Vesuna, K. Evans, C. Liu, C. Ramakrishnan, J. Liu, et al. Threedimensional intact-tissue sequencing of single-cell transcriptional states. Science, 361(6400), 2018.

[63] M. D. Wilkinson, M. Dumontier, I. J. Aalbersberg, G. Appleton, M. Axton, A. Baak, N. Blomberg, J.-W. Boiten, L. B. da Silva Santos, P. E. Bourne, et al. The fair guiding principles for scientific data management and stewardship. Scientific data, 3(1):1–9, 2016.

[64] F. A. Wolf, P. Angerer, and F. J. Theis. Scanpy: large-scale single-cell gene expression data analysis. Genome biology, 19(1):1–5, 2018.

